# Detailed spectrographic analysis of rat ultrasonic vocalizations emitted during the acoustic startle response test

**DOI:** 10.1101/2023.02.24.529853

**Authors:** Emilie Bartsoen, Markus Wöhr

**Affiliations:** KU Leuven, Faculty of Psychology and Educational Sciences, Research Unit Brain and Cognition, Laboratory of Biological Psychology, Social and Affective Neuroscience Research Group, B-3000, Leuven, Belgium; KU Leuven, Leuven Brain Institute, B-3000, Leuven, Belgium; Philipps-University of Marburg, Faculty of Psychology, Experimental and Biological Psychology, Behavioral Neuroscience, D-35032, Marburg, Germany; Philipps-University of Marburg, Center for Mind, Brain and Behavior, D-35032 Marburg, Germany

**Keywords:** ultrasonic communication, ultrasound, vocalizations, acoustic startle reflex, emotional state, habituation type

## Abstract

Rats emit ultrasonic vocalizations (USV). During aversive situations, rats emit 22-kHz USV, which are considered “alarm calls” and supposed to reflect a negative affective state of the sender. During appetitive situations, rats emit 50-kHz USV, which are believed to reflect a positive affective state. Here, we recorded USV emission in adult male rats during the acoustic startle response test. Our results indicate varied USV emission in both the 22- and 50-kHz USV ranges. Enhanced startle responses were observed in rats with a predominant 22-kHz call profile, supporting the notion that 22-kHz USV emission is associated with a negative affective state.

---

Depending on the emotional valence of a situation, rats emit different types of ultrasonic vocalizations (USV) [1,2]. 22-kHz USV have a low peak frequency (<32kHz) and a long duration (>300ms) [3–5]. They are emitted in aversive situations (e.g., the acoustic startle response test), and are therefore seen as an expression of a negative affective state [1,2]. They serve as “alarm calls” and cause a fear-like response in the receiver (e.g., behavioral inhibition, amygdala activation) [6,7]. In contrast, 50-kHz USV have a high peak frequency (>32kHz), are typically short (<300ms) and can be divided into categories according to their level of frequency modulation (e.g., flat, trill) [2–5]. They are mainly emitted during appetitive situations (e.g., rough-and-tumble play) and evoke reward-associated responses (e.g., approach behavior, dopamine release) [8,9]. 50-kHz USV are believed to be an index of the rat’s appetitive affective state and serve to restore social contact [1,2].

The acoustic startle response is a contraction of the body muscles in response to loud noise bursts. Its amplitude depends on the affective state of the subject. In rats, this was shown in several studies in which a stimulus presented during the ASR test led to a decrease or increase of the startle amplitude depending on its association with reward or punishment, respectively [10–14]. Despite its importance in affective neuroscience, the measurement protocol for the ASR varies widely from study to study. For example, it is not uniform how rats are habituated to the startle equipment before the test [e.g., 13,14], and in some studies no prior habituation was conducted [e.g., 11,12]. Prior habituation may impact the affective state and thus the startle amplitude of the rat, making comparison of studies difficult.

A few studies measured USV emission during the ASR test, reporting only 22-kHz USV [15–18]. However, such studies were conducted about 30 years ago and at that time most USV recordings were still technically limited to narrow frequency bandwidths, likely resulting in an underrepresentation of 50-kHz USV [19]. Therefore, it remains to be determined whether the call profile is more versatile during the ASR test when all frequencies are detected. To fill this gap, we conducted broadband USV recordings on rats during the ASR test and analyzed them in a detailed manner. Rats were divided into three habituation types, and their effect on USV emission and startle amplitude was assessed. Finally, since both USV emission and startle amplitude are believed to be overt expressions of the affective state, it was examined whether the ASR differed between rats depending on their calling behavior.

Thirty-six male Sprague-Dawley rats (Charles-River, France; weight: 245-300g) were used. The rats were housed with four in polycarbonate Macrolon type IV cages (33×53×19cm; plus high stainless steel lids) with Safe^®^ flake bedding. Climate conditions were controlled (light on: 7a.m. to 7p.m., temperature: 21-24°C, humidity: 30-40%) and food and water were provided *ad libitum*. Experiments were performed in accordance with the European Communities Council Directives and permitted by the local animal ethics committee (P179/2021).

The ASR was measured using the SR-LAB startle response system (San Diego Instruments, USA). The equipment consisted of a cylindrical animal enclosure (⌀: ~9cm, L: ~20cm) placed in a soundproof isolation chamber (~33×33×48cm). The animal enclosure was mounted on a Plexiglas base (~20×13cm) with a piezoelectric accelerometer underneath. The accelerometer detected the rat’s startle response for 200 ms after the onset of the noise burst at a frequency of 200 Hz. The startle responses were processed by the SR-LAB software and digitized as data points. Noise bursts of 7 different intensities (78, 85, 92, 99, 106, 113, 120dB; duration: 40ms) were administered through a loudspeaker mounted ~24 cm above the animal. The intensity of the noise bursts was measured with a 47730 digital sound level meter (Extech Instruments, USA) directly below the speaker, and monitored via the SR-LAB software. The baseline response of the accelerometer was calibrated using the SR-LAB standardization unit.

Prior to the experiment, all rats were handled for 3 consecutive days according to a standardized protocol (5 min/day). To test the effect of habituation on the ASR, rats were randomly divided into three groups (N=12 per group) on the first day: 1) Habituation group: 5 min habituation to the startle equipment, 2) Habituation + noise bursts group: 5 min habituation to the startle equipment plus four 120 dB noise bursts (inter-stimulus interval: 30s), or 3) No habituation group: the rats remained in the home cage. On the second day of the experiment, all rats were subjected to the ASR test (Figure 1A). During this test, all 7 noise bursts intensities were presented in blocks (10 blocks with 70 noise bursts in total) after a 5-minute acclimatization period. The inter-stimulus interval varied between 5-30 s (mean: 17.35s) and the sequence of noise burst intensities was randomized between blocks. Continuous background noise (60dB) and moderate internal light were present during the test. The mean peak startle response was calculated per noise burst intensity for each rat.

**Figure 1:**
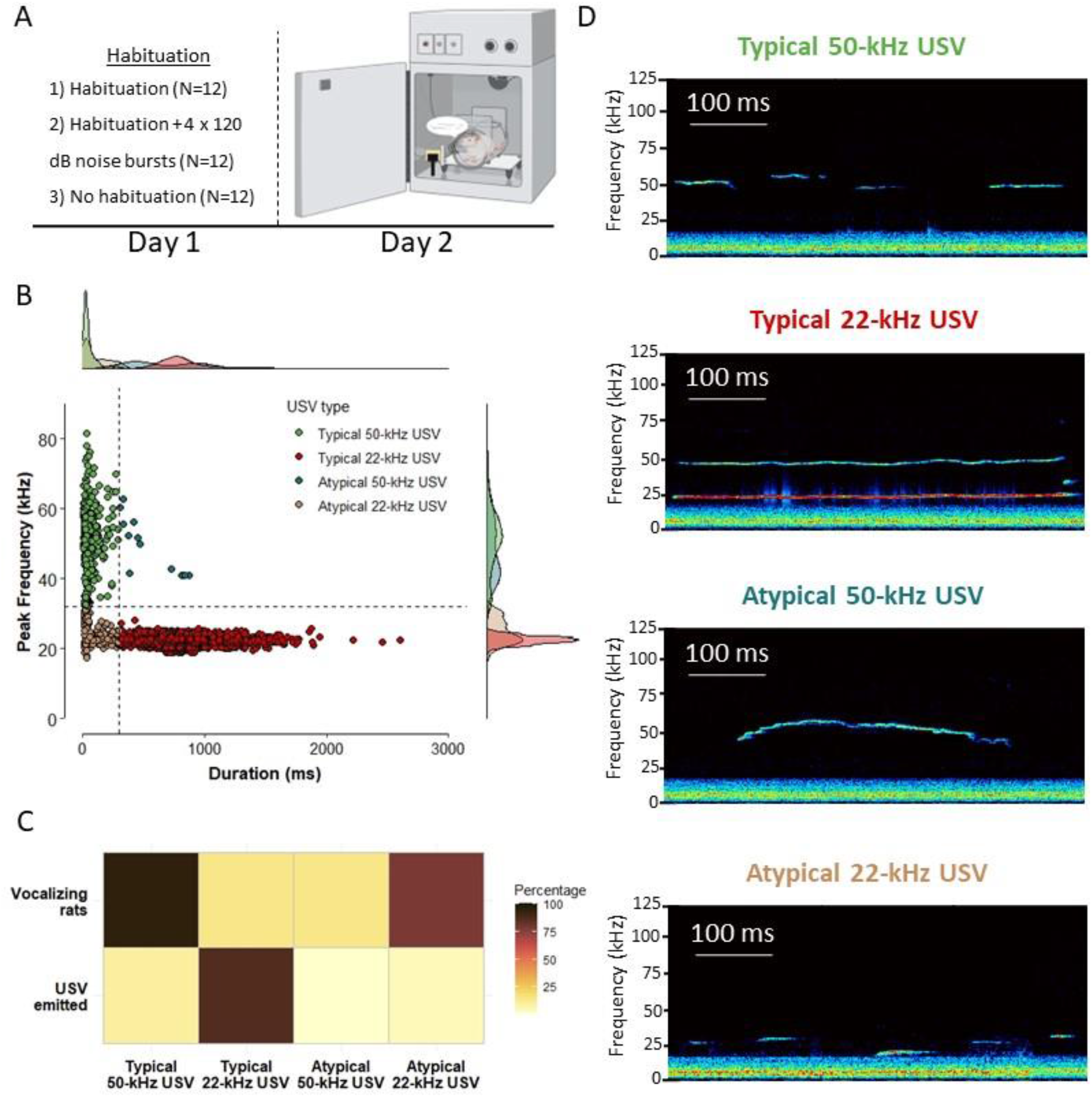
A) Overview of the experimental design. Rats were divided into habituation groups on the first day and exposed to the ASR test on the second day. B) Scatterplot showing all emitted USV plotted as a function of duration (ms) and peak frequency (kHz). Dashed lines indicate the cutoff values used to define USV types. The densities for each USV type along the axes for duration and peak frequency are shown at the sides. C) Percentage of rats that emitted each USV type (top row) and percentage per USV type of all emitted USV (bottom row). D) Exemplary spectrograms of each USV type. Please note that the exemplary spectrogram of a typical 22-kHz USV includes a prominent overtone.

During the ASR test, USV were recorded with an UltraSoundGate condenser microphone (CM16, Avisoft Bioacoustics, Germany) mounted in the right corner of the soundproof isolation chamber ~4 cm away from the animal. The microphone was connected to a personal computer via an Avisoft Ultrasound Gate 416 USB Audio Device, where the USV were recorded with Avisoft Recorder 4.3.01 (sampling rate: 250,000Hz; format: 16 bit). Using Avisoft SASLabPro 5.3.01 Software, high-resolution spectrograms (frequency resolution: 0.488kHz, time resolution: 0.512ms) were generated by applying a Fast-Fourier transform (512 FFT length, 100% frame, Hamming window, 75% time window overlap). USV were manually labelled and the software automatically calculated call number, peak frequency (kHz), duration (ms) and frequency modulation (kHz, i.e., the difference between the highest and lowest frequency of the call). Based on cut-off values used in previous studies [3–5], the USV were divided into categories according to duration and peak frequency (Figure 1B): typical 22-kHz USV (duration >300ms, peak frequency <32kHz), atypical 22-kHz USV (duration ≤300ms, peak frequency <32kHz), typical 50-kHz USV (duration ≤300ms, peak frequency ≥32kHz), or atypical 50-kHz USV (duration >300ms, peak frequency ≥32kHz; Figure 1D).

Statistical analyses were performed in RStudio 4.1.2. To examine the effect of habituation group and USV type on the ASR, mixed ANOVA analyses were used with noise burst intensity as within-subject variable. If significant, pairwise comparisons with Bonferroni correction were applied (Bonferroni-adjusted *p*-values are denoted as *p_adj_*). Chi-squared tests were used to compare the proportion of each USV type between habituation groups.

Rats were highly vocal during the ASR test and produced versatile calls. Almost all rats (97.22 %) emitted at least one but mostly many more USV, resulting in a total of 3896 calls. The majority of calls emitted during the ASR test were typical 22-kHz USV (86.19%), followed by typical 50-kHz USV (9.39%), atypical 22-kHz USV (4.06%), and atypical 50-kHz USV (0.36%). Notably, although more typical 22-kHz USV were observed, they were emitted by fewer rats (13.89%, N=5 rats) than typical 50-kHz USV (94.44%, N=34 rats) and atypical 22-kHz USV (77.78%, N=28 rats) (Figure 1C). More specifically, if rats emitted typical 22-kHz USV, they generally did so continuously throughout the test session, while the other USV types were emitted by more rats, but not continuously. The atypical 50-kHz USV were rare and emitted by only a few rats (13.89%, N=5 rats).

The typical 22-kHz USV were long (762.56±4.37ms) and had a peak frequency of 22.48±0.02 kHz. They were characterized by a flat call profile (mean frequency-modulation ≤5kHz), where oftentimes a frequency step could be seen at the beginning and/or end of the call. The atypical 22-kHz USV, on the other hand, were also flat in nature (mean frequency-modulation ≤5kHz) but had a shorter duration (100.82±7.16ms) and a slightly higher peak frequency (24.92±0.27kHz) compared to typical 22-kHz USV. The typical 50-kHz USV were short (55.63±2.84ms), with a high peak frequency (51.13±0.48kHz). Strikingly, these calls showed minimal frequency-modulation (≤5kHz), to a similar extent as both 22-kHz USV categories. This was in contrast with the atypical 50-kHz USV that were highly frequency-modulated (>5kHz) and characterized by long durations (568.46±61.20ms). Notably, their peak frequency (48.44±2.16kHz) was lower compared to typical 50-kHz USV. An overview of the call parameters per category is displayed in Table 1.

**Table 1:** Call parameters per USV type

|  | Typical<br>22-kHz USV | Atypical<br>22-kHz USV | Typical<br>50-kHz USV | Atypical<br>50-kHz USV |
| --- | --- | --- | --- | --- |
| Total USV (%; $N=3896$) | 86.19 | 4.06 | 9.39 | 0.36 |
| Rats (%; $N=36$) | 13.89 | 77.78 | 94.44 | 13.89 |
| Duration (ms) | $762.56 \pm 4.37$ | $100.82 \pm 7.16$ | $55.63 \pm 2.84$ | $568.46 \pm 61.20$ |
| Peak frequency (kHz) | $22.48 \pm 0.02$ | $24.92 \pm 0.27$ | $51.13 \pm 0.48$ | $48.44 \pm 2.16$ |
| Frequency-modulation (kHz) | $4.99 \pm 0.07$ | $4.04 \pm 0.28$ | $5.00 \pm 0.31$ | $13.44 \pm 2.21$ |

Data is presented as mean ± SE

Next, the effect of habituation on ASR and USV emission across different noise burst intensities was studied. The startle amplitude did not differ between habituation groups across noise burst intensities (main effect: F(2,33)=1.06, *p*=0.375; interaction effect: F(2.85,47.06)=0.91, *p*=0.440; Figure 2A). Of note, the measurement of the ASR is highly dependent on body weight, however, as weight did not differ between habituation groups (F(2,33)=0.50, *p*=0.613), it was not included as a covariate in the analysis. Habituation did also not affect USV emission during the ASR test. The percentage of rats emitting USV did not differ between habituation groups (X^2^(2)<0.01 *p*=0.998), even when distinguishing between different USV types (Figure 2B; Typical 22-kHz USV: X^2^(2)=0.43, *p*=0.805; Typical 50-kHz USV: X^2^(2)=0.01, *p*=0.805; Atypical 22-kHz USV: X^2^(2)=0.01, *p*=0.997; Atypical 50-kHz USV: X^2^(2)=0.23, *p*=0.890). Due to the absence of an effect on both ASR and USV emission, habituation group was not considered for further analyses.

**Figure 2:**
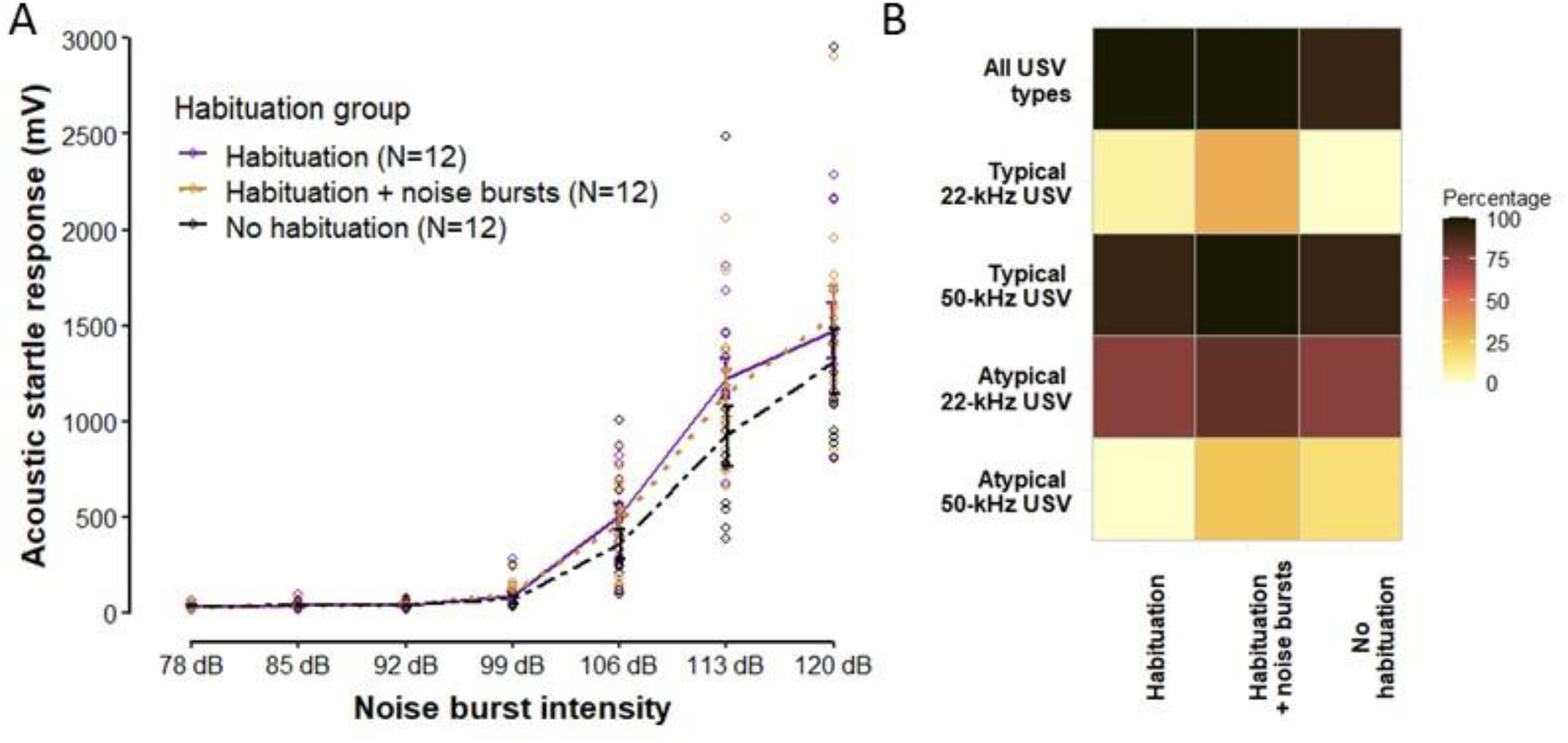
A) Line graph showing the effect of habituation on the acoustic startle response for different noise burst intensities. For each noise burst intensity, the mean±SE for each habituation group is plotted. Points represent the mean acoustic startle response of individual rats. B) Proportion of rats emitting USV per habituation group, either total USV emission or by USV type.

To examine whether inter-individual differences in USV emission are associated with the startle amplitude, only the typical 22-kHz USV and typical 50-kHz USV were included for feasibility reasons. Consequently, based on their call profile, rats were classified into 1) the “No USV” group, if they produced 4 or fewer calls, 2) the “22-kHz USV” group, if more than 50% of their calls were typical 22-kHz USV, or 3) the “50-kHz USV” group, if more than 50% were typical 50-kHz USV. This resulted in N=10, N=5, and N=21 rats in the “No USV”, “22-kHz USV”, and “50-kHz USV” groups, respectively. When comparing the time spent calling, prominent differences were seen between groups (“No USV” group: 0.00±0.00s 22-kHz USV and 0.07±0.02s 50-kHz USV; “22-kHz USV” group: 512.14±138.08s 22-kHz USV and 0.20±0.09s 50-kHz USV; “50-kHz USV” group: 0.00±0.00s 22-kHz USV and 0.89±0.21s 50-kHz USV).

While the startle amplitude increased with noise burst intensity irrespective of inter-individual differences in USV emission (F(6,198)=181.20, *p*<0.001), the steepness of this increase was clearly dependent on the USV group (F(12,198)=2.08, *p*=0.020; Figure 3A). This interaction effect was most prominent at the loudest noise burst (120dB), where rats in the “22-kHz USV” group displayed a stronger ASR compared to the “50-kHz USV” group (*p*=0.028) and marginally higher compared to the “No USV” group (*p*=0.054) (The pairwise comparisons did not survive Bonferroni correction). Importantly, rats in the “No USV” group had a lower body weight than rats in both the “22-kHz USV” (*p_adj_*=0.005) and “50-kHz USV” (*p_adj_*<0.001) groups (F(2,33)=9.65, *p*<0.001; Figure 3B), yet the enhanced startle amplitude in the “22-kHz USV” group remained significant after including weight as a covariate (F(12,192)=1.90, *p*=0.036; after correcting for sphericity: F(2.83,45.24)=1.90, *p*=0.146).

**Figure 3:**
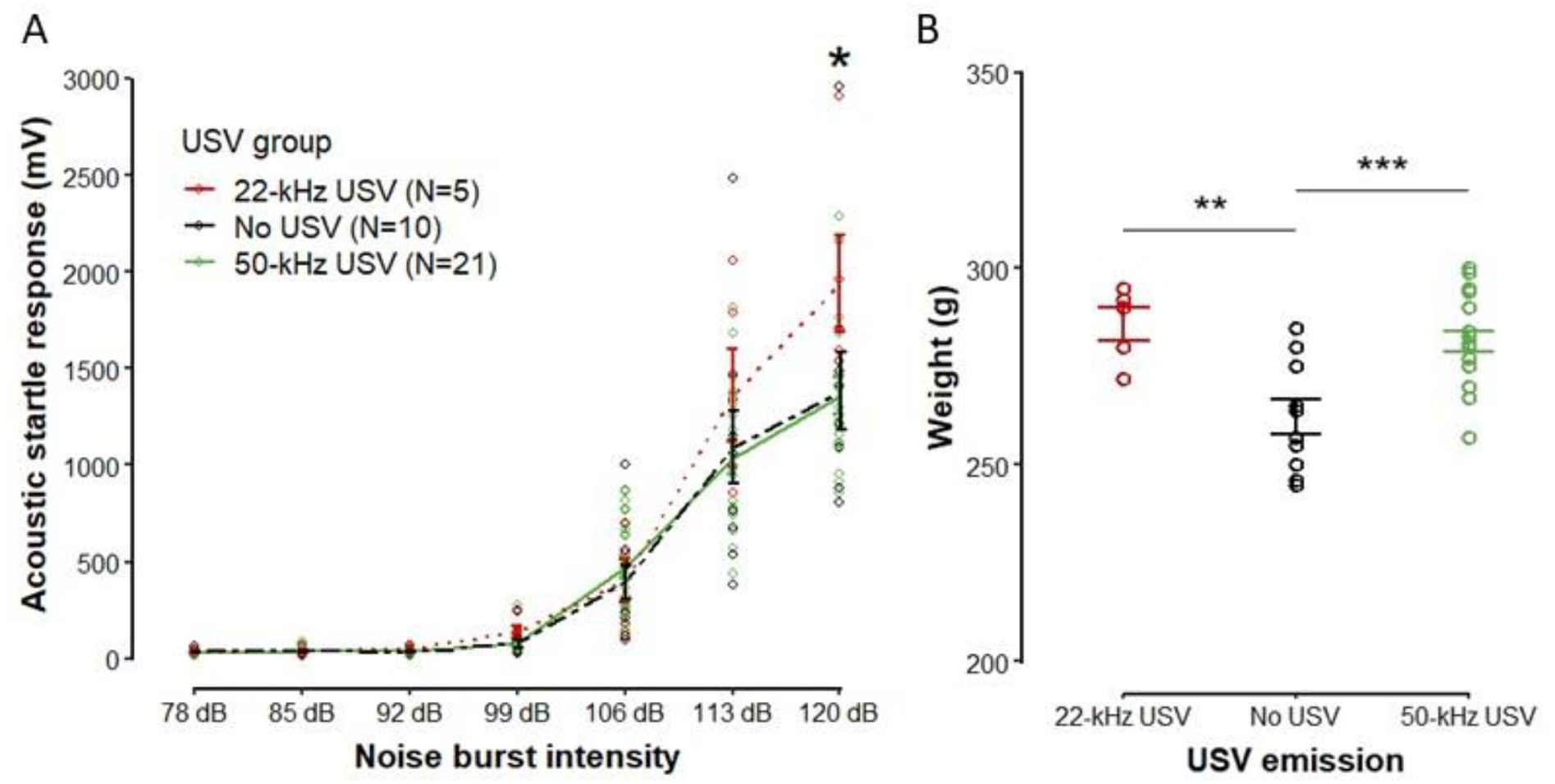
A) Line graph showing the acoustic startle response with increasing noise burst intensities separately for each USV group. B) Body weight per USV group. For both graphs, bars visualize mean±SE and points represent individual rats. \**p*<0.05, \*\**p*<0.01, \*\*\**p*<0.001

In the present study, rats were subjected to the ASR test while recording USV emission. The main findings were threefold: 1) Numerous USV of variable durations and frequencies were observed. Typical 50-kHz USV were most often emitted, followed by atypical 22-kHz USV, typical 22-kHz USV and atypical 50-kHz USV. 2) Whether rats were habituated to the startle equipment, and whether this habituation was combined with noise bursts, did not affect either startle amplitude or USV emission. 3) Rats that emitted mainly typical 22-kHz USV had an enhanced startle response at the loudest noise burst.

Numerous studies have reported large inter-individual variability among rats in their call profile [20,21]. Also in the present study, variability was observed between rats in their call rate and the type of calls that were emitted. Most rats emitted short and flat 50-kHz USV, while long frequency-modulated 50-kHz USV and long flat 22-kHz USV were emitted by the fewest rats. Interestingly, the few studies that have focused on USV emission during the ASR test only report observations of long typical 22-kHz USV and none in the 50-kHz range [15–18]. However, their USV recordings were restricted to narrow frequency bandwidths due to technical limitations, which might have under-sampled 50-kHz USV. Nonetheless, when comparing the percentage of 22-kHz USV emitting rats, there is a notable difference. Previous studies report ASR-induced 22-kHz USV emission in 40-70% of their rats. The emission typically started after the first few noise bursts and was continued throughout the test. In our study, a large percentage of rats did occasionally emit atypical 22-kHz USV, but continuous emission of long typical 22-kHz USV was only observed in 5 rats (13.89%). This discrepancy in call rate is most likely due to differences in the ASR protocol. We used a wide range of noise burst intensities, where loud noise bursts (i.e. 106, 113, 120 dB) were alternated with lower intensities (i.e. 78, 85, 92, 99 dB). In contrast, previous studies [15–18] only presented louder noise bursts (i.e. 105, 110, 115 dB). This might have induced a more anxious state in these rats, with an increase in 22-kHz USV emission as a consequence.

The magnitude of the ASR is a validated tool in both rats and humans to assess emotional reactivity [22]. By presenting a conditioned stimulus that was associated with an aversive or appetitive event, the amplitude of the ASR is enhanced or reduced, respectively. The emotional state is thus directly linked to the ASR amplitude, which is enhanced by a negative affective state (e.g., fear) and reduced by a positive affective state (e.g., reward anticipation) [10–13]. In the present study, rats that emitted primarily 22-kHz USV had a higher startle response to 120 dB noise bursts compared to rats that emitted primarily 50-kHz USV or were silent for the entire test. Importantly, this effect was not confounded by higher weight in the 22-kHz USV emitting rats compared to the silent rats, as the effect remained significant after correcting for the weight difference. Moreover, the largest difference in ASR amplitude was found between 22-kHz and 50-kHz USV emitting rats, which did not differ significantly in weight. In line with our findings, Vivian et al. (1994) [18] found that rats experiencing drug withdrawal exhibited increased 22-kHz USV emission that was accompanied by an increased ASR (significant at 115 dB noise bursts). 22-kHz USV emission is ethologically relevant for rats and they are mainly emitted in aversive contexts, for example, when they receive shocks or smell a predator [1,2]. These calls are therefore hypothesized to be an index of a negative affective state, similar to fear [1,2]. Our findings support this hypothesis, where rats most likely experienced fear during the ASR test, which was expressed by emitting 22-kHz USV, resulting in an enhanced startle response.

It was surprising that 50-kHz USV were emitted during the ASR test, which is an aversive context for rats. In addition, it was unexpected that rats emitting 50-kHz USV did not display a reduced ASR. For a correct interpretation of these results, it is important that the observed 50-kHz USV were flat in nature, and that these calls have been shown to be functionally different from frequency-modulated 50-kHz USV. Frequency-modulated 50-kHz USV are emitted in appetitive contexts (e.g., rough-and-tumble play, tickling, mating, psychostimulant drugs), while flat calls are emitted in mildly stressful contexts (e.g., open field test, aggressive encounters) [1,2,20,21,23]. Rats also specifically perceive frequency-modulated calls as rewarding, as they actively self-administer or approach playback of frequency-modulated, but not flat 50-kHz USV [23]. Therefore, it was suggested that frequency-modulated 50-kHz USV reflect a positive affective state, while flat 50-kHz USV serve as non-affective contact calls to initiate social contact [21,23]. This was supported by studies, where rats exposed to brief social isolation emitted flat 50-kHz USV that decreased over time [20,23]. Also in the present study, rats underwent the ASR test in isolation, and therefore the emitted flat 50-kHz USV presumably served to restore contact with their cage mates. The fact that these calls have a social directing function, rather than an indicator of a positive affective state, would explain why no reduction in their startle response was observed.

In sum, we demonstrated by detailed spectrographic analysis a versatile range of emitted USV during the ASR test with respect to duration and frequency. Our results also highlight the ethological significance of the different USV types by demonstrating ASR potentiation associated with 22-kHz USV emission, supporting the notion that 22-kHz USV emission is associated with a negative affective state. This highlights the power of the ASR as a window into the affective state of rats.

## Acknowledgements

The authors wish to thank Robin Vloeberghs and Nathan Van Humbeeck for their helpful suggestions for the statistical analyses. This project was supported by the Fonds Wetenschappelijk Onderzoek – Vlaanderen (FWO; Research Foundation – Flanders) through a senior project to M.W. (G0C0522N) and the Interne Fondsen KU Leuven (Internal Funds KU Leuven) through a BOFZAP Starting Grant to M.W. (PXF-E0120-STG/20/062).

